# Augmenting the Signal Peptide of the Ag43 Autotransporter for the improved heterologous display of sfGFP using Fluorescence-Activated Cell Sorting (FACs)-assisted natural selection

**DOI:** 10.1101/2022.07.28.501931

**Authors:** Darius Wen-Shuo Koh, Jian-Hua Tay, Samuel Ken-En Gan

**Affiliations:** Antibody & Product Development Lab, Agency for Science, Technology, and Research (A*STAR), Singapore; James Cook University, Singapore, Singapore; Zhejiang Bioinformatics International Science and Technology Cooperation Center, Wenzhou-Kean University, Wenzhou, Zhejiang Province, China; Wenzhou Municipal Key Lab of Applied Biomedical and Biopharmaceutical Informatics, Wenzhou-Kean University, Wenzhou, Zhejiang Province, China

**Author notes:** These authors contributed equally (Darius Wen-Shuo Koh, Jian-Hua Tay).

**Keywords:** Single cell screening, Protein surface display, Engineering signal peptides, Ag43 Autotransporter, error-prone PCR

## Abstract

Protein display, secretion and export in prokaryotes are essential for utilizing microbial systems as engineered living materials for medicines, biocatalysts, and protein factories. To select for improved signal peptides for *Escherichia coli* protein display, we utilized error-prone polymerase chain reaction (epPCR) coupled with single-cell sorting and microplate titer to generate, select, and detect improved Ag43 signal peptides. Through three rounds of mutagenesis and selection using green fluorescence from the 56 kDa sfGFP-beta-lactamase, we isolated clones that increased surface display from 1.4 to 3 folds as detected by the microplate plate-reader and native SDS-PAGE assays. To establish that the protein was displayed extracellularly, we trypsinised the bacterial cells to release the surface displayed proteins for analysis. This workflow demonstrated a fast and high-throughput method leveraging on epPCR and single-cell sorting to rapidly augment bacterial surface display, a method that could be applied to other bacterial proteins.

## Introduction

Augmenting protein display and secretion by bacteria is desirable for many applications such as the development of whole cell catalysts (Hörnström et al., 2019) in industrial biotechnology and the display of vaccine subunits (Schroeder et al., 2011) as living medicines. Many approaches from the optimization of the ribosome binding site, promoter, and codon usage to the screening of different natural signal peptides or different autotransporter β-barrel domains to optimize the autotransporter mediated display of recombinant secreted proteins (Burdette et al., 2021; Gustavsson et al., 2011a) demonstrate the interest in this area.

To enable the programming of cells to fabricate engineered living materials (ELMs) (Duraj-Thatte et al., 2021; Glass and Riedel-Kruse, 2018; Jin and Riedel-Kruse, 2018; Widmaier et al., 2009), deliver protein therapeutics, (Cui et al., 2021; Ho et al., 2018; Veiga et al., 2003; Wu et al., 2013) test mutant protein variants faster (Alessa et al., 2019; Daugherty et al., 2000; Ling et al., 2022, 2018; Starr et al., 2020; Su et al., 2021, 2018; Warszawski et al., 2019), optimize bioprocess parameters (Heng et al., 2022; Ling et al., 2020), perform biochemical reactions(Lindroos et al., 2019; Muñoz-Gutiérrez et al., 2014) and construct biosensors (Kylilis et al., 2019; VanArsdale et al., 2020), require the production, display or secretion of proteins in both eukaryotes and prokaryotes. *Escherichia coli* (*E. coli*) remains one of the best-characterized model organisms for engineering to secrete proteins into the extracellular environment (Kleiner-Grote et al., 2018). Given that proteins can be either displayed on the surface of cells or released into the medium (Ahan et al., 2019; Azam et al., 2016), there is the opportunity to harvest these proteins without a laborious lysis step that could involve contamination of endogenous intracellular proteins. With extracellular expression reported to circumvent cytosolic aggregation and proteolysis of proteins, higher titres of untruncated protein-based polymeric materials (Azam et al., 2016) could be harvested from surface displayed proteins.

The secretion and protein export pathways of *E. coli* are divided into Type I, II, III, IV and V (reviewed in (Kleiner-Grote et al., 2018; Meuskens et al., 2019)) pathways. Many attempts were made to engineer these pathways for heterologous protein production and export, mainly targeting the Type I system, where attempts to modify it usually encompass mutagenesis of secretion carrier proteins (Gonzalez-Perez et al., 2021), leader peptides(DeLisa et al., 2002; Selas Castiñeiras et al., 2018) and chaperones (Haitjema et al., 2014; Natarajan et al., 2017; Taw et al., 2021) in the Sec/Tat pathways followed by the selection and screening systems that were based on high throughput plate screening or antibiotic survival selection assays.

The natural Ag43 autotransporter system usually consists of a signal peptide (to mediate timing and targeting by chaperones), an N-proximal passenger domain (α^43^), auto chaperone domain, and a C-terminal Ag43 β-barrel domain (β^43^) (Ahan et al., 2019; Ramesh et al., 2012; van der Woude and Henderson, 2008). It is abundantly displayed on the bacteria and belongs to the Type V, subtype A, secretion system, relying on the Sec chaperone system (Meuskens et al., 2019; Ramesh et al., 2012; van der Woude and Henderson, 2008). The Ag43 autotransporter enables protein secretion into the media through a proteolytic site in α^43^ (Ahan et al., 2019; Ramesh et al., 2012; van der Woude and Henderson, 2008), where the absence of this domain would allow for heterologous proteins to be more reliably displayed (Ahan et al., 2019; Ramesh et al., 2012). One advantage of the Ag43 system is its ability to display a vast repertoire of disulphide-containing proteins as determined by single-cell flow cytometry (Kjærgaard et al., 2002; Ramesh et al., 2012). With the many uses of the Ag43 autotransporter system, such as engineering the specific gut colonization of probiotic *E. coli* Nissle 1917 strain to display adhesive proteins to alleviate dextran sulfate sodium-induced colitis in murine models (Cui et al., 2021), and thermostable β-glucosidase display for whole-cell biocatalyst transformation of cellulose into simple sugars (Muñoz-Gutiérrez et al., 2014), there remain many applications. Examples in the heterologous expression of proteins are that they could also be toggled between display and secretion modes with conditional expression of tobacco etch protein proteases(Ahan et al., 2019) and as ELMs in allowing for microbial optogenetic lithography of complex spatial biofilm patterns (Jin and Riedel-Kruse, 2018).

One advantage of engineering the Ag43 system is that the protein of interest can be physically linked to the cell, thus allowing for fluorescence-activated cell sorting (FACs) – a powerful multiparametric and (semi) quantitative tool (Beal, 2015; Georgiou, 2001) widely used in bio-foundries (Holowko et al., 2020; Pham et al., 2017) to develop microbes resistant to low pHs (Pham et al., 2017), superfast enzymes (Griffiths and Tawfik, 2003), LacI (Taylor et al., 2016) and Ribosensors (Townshend et al., 2015) with expanded ligand responsiveness which could not otherwise exist in nature.

To engineer the Ag43 system for a more productive display and export of a moderately sized passenger protein or sfGFP-beta-lactamase fusion (56 kDa) beyond the cell wall, we utilized error-prone polymerase chain reaction (epPCR) with FACs in a proof-of-concept experiment for the selection of augmenting protein export in *E. coli* with validation and analysis using plate reader high-throughput measurements. Although the switching of natural signal peptides to optimize signal peptides of autotransporters was previously made (Gustavsson et al., 2011b; Ramesh et al., 2012), the methodology using epPCR to mutate signal peptides of autotransporters with single cell selections reported here is to our knowledge, previously reported.

## Materials and Methods

### An overall flow of the methodology is as shown in Figure 1

**Figure 1.**
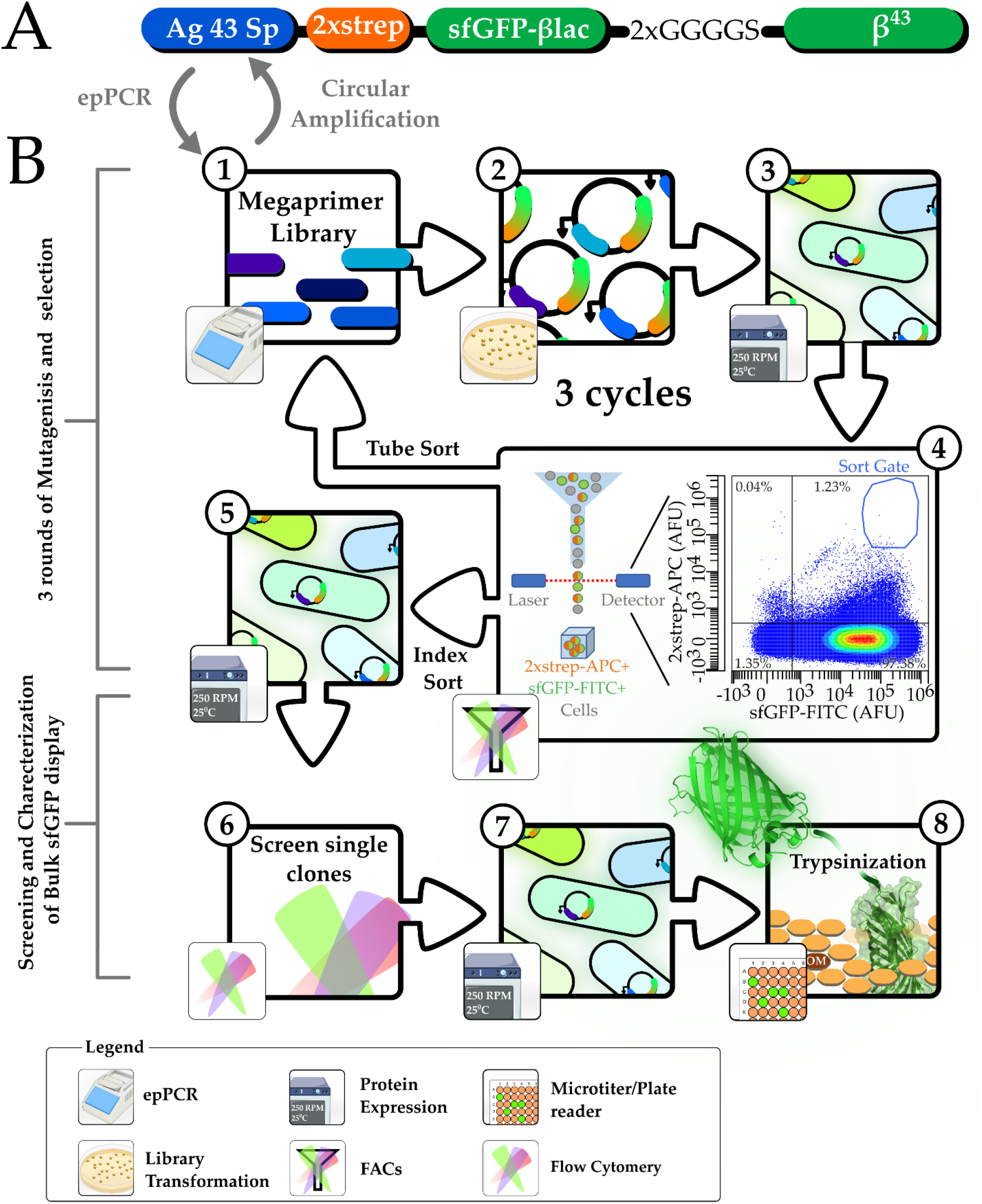
Schematic of the workflow to generate and select for Ag43 signal peptide (Ag43 Sp) mutant sequences for augmented display. (**A**) Illustration of the critical features in the Ag43 Autotransporter system: Ag43 signal peptide (Ag43 Sp), twin-strep tag-II(2xstrep), sfGFP, β-lactamase (βlac), 2x GGGGS linker and β-barrel domain(β^43^) which also tethered the protein onto the outer membrane. (**B**) The workflow to generate the Ag43 signal peptide mutations in this study; steps are labelled in circular labels. (**Step 1-4 mutagenesis and selection, repeated for three cycles**). Step 1, Megaprimer libraries were generated with error-prone PCR (epPCR) to create megaprimers used for circular amplification (using wildtype plasmid pCDF_Ag43sp_2xstrepSfGFPβLac as template) to generate mutant signal peptides. Steps 2 and 3, transformation and harvesting of mutant colonies for subsequent expression with 0.5 mM IPTG at 25°C, 250 rpm for 5 to 6 hours. In step 4, mutant Ag43 bacteria libraries were sorted with FACs. (**Steps 5-8, screening, and characterization of isolated index sorted clones**) steps 5-6, 43 clones (index sorted into microwell plates) were expressed and screened with FACs. In steps 7-8, 3 of the best clones (with the best Flow-Cytometric profiles) were selected and expressed in 50 ml falcon tubes and trypsinization to release displayed sfGFP on the cell surface. Trypsinized cell supernatants were scanned for sfGFP signals with a plate-reader and subjected to native SDS-PAGE for analysis.

### Molecular cloning

To design the Ag43 autotransporter display system, a codon-optimized sfGFP sequence was flanked by the native nucleotide sequence (NCBI GeneID: 94650) for the N terminal Ag43 signal peptide, twin strep-tag II (2xstrep) and C terminal β^43^ domain as performed by Ramesh et al., (2012) and illustrated in Figure 1A. The designed construct was gene synthesized and cloned into the pCDF plasmid backbone using the Ncol and XhoI restriction sites added by Genscript with an alanine codon added after the first methionine in the signal peptide to ensure in-frame with the Ncol restriction site.

To create the final construct used in this study(pCDF_Ag43sp_2xstrepSfGFPβLac, Figure 1A), the β-Lactamase gene was fused to the 3’ end of the sfGFP protein to make a moderately sized 56 kDa model passenger protein by first mutagenizing pCDF_Ag43sp_2xstrepSfGFPβLac by primers: Ag43LinkBlac_F, 5’-ATG GAC GAA TTA TAT AAA GGT ACC GAG GAT CTG TAC TTT CAG AGC GGT GGA GGT GGT TCT GGT GGT GGA GGT-3’; and Ag43LinkBlac_R, 5’- ACC TCC ACC ACC AGA ACC ACC TCC ACC GCT CTG AAA GTA CAG ATC CTC GGT ACC TTT ATA TAA TTC GTC CAT-3’) using the Agilent QuikChange SDM kit following the manufacturer’s instructions (Agilent, Singapore, Singapore; Cat: STR_210519). This was followed by amplifying the backbone of the plasmid using primer set Ag43BBGib_F, 5’- GGT ACC TTT ATA TAA TTC GTC CAT ACC ATG CGT −3’; and Ag43BBGib_R, 5’- GCT GCT GCA AGC GAG GAT CTG TAC TTT CAG AGC GG -’3 with the Platinum SuperFi II Green PCR master mix (ThermoFisher, Singapore; Cat: 12369010). The polymerase chain reaction (PCR) profile is as follows: initial denaturation at 95°C for 120 seconds, followed by 15 cycles of touch-down PCR (denaturation at 95°C for 30 seconds, annealing at 72 °C for 30 seconds with auto-delta every cycle set as −1 °C after the first cycle) and extension at 72°C for 180 seconds and 25 cycles of conventional PCR (done similarly as per touch-down PCR except for the annealing temperature fixed at 60°C). β-Lactamase gene (amplified from Addgene plasmid RRID: #52656; using primer set Ag43BlacGib_F, 5’- CAG ATC CTC GCT TGC AGC AGC CCA ATG CTT AAT CAG TGA GGC ACC −3’; and Ag43BlacGib_R, 5’- TGG TAT GGA CGA ATT ATA TAA AGG TAC CGG CGG AGG CTC CCA CCC AGA AAC GCT GGT GAA AGT AAA AGA T −3’) were amplified in the same manner except that the extension time was set at 25 seconds. The gel-extracted plasmid backbone and insert were assembled with 2x NEB HiFi DNA assembly kit following the manufacturer’s instructions.

Plasmids were prepped as previously described (Poh and Gan, 2014) using *Escherichia coli* BL21 (DE3) previously described in (Chan et al., 2013). DNA quantifications were performed with the Qubit DS HS or BR DNA kit (Invitrogen™, Singapore) following the manufacturer’s instructions. High throughput optical density (OD) normalization steps were performed with the OT2 (OpenTrons, USA) liquid handling automation system using custom python scripts. For pCDF based plasmids, 200 μg/ml of spectinomycin was used. All centrifugation procedures were performed at 8000 rpm for 5 minutes or 4000 rpm for 1 hour in 1.5 ml microfuge tubes or 2ml Deepwell (NEST, Cat: 503501) plates for low and high throughput procedures, respectively.

### Mutant library construction

Plasmids isolated from the gene synthesized clones and FAC sorted BL21 (DE3) *E. coli* cells were subjected to high fidelity Q5 Polymerase (NEB, Singapore; Cat: M0491L) PCR to generate 200 bp templates of the Ag43 signal peptide (Figure 1A). To generate the mutant Ag43 signal peptide library, 2.5 ng of the template per 100 μl were then subjected to epPCR (with primer set Ag43epPCR_F, 5’- CTG TAG AAA TAA TTT TGT TTA ACT TTA ATA AGG AGA TAT ACC ATG −3’; and Ag43epPCR_R, 5’- CTT TTT CAA ATT GGG GGT GTG ACC ATG CAG CGC TAG CCG CAG C −3’) using the GeneMorph II epPCR kit (Agilent, Singapore, Cat: STR_200550). The epPCR profile is as follows: initial denaturation at 95°C for 120 seconds, followed by 15 cycles of touch-down PCR (denaturation at 95°C for 30 seconds, annealing at 72 °C for 30 seconds with auto-delta every cycle set as −1 °C after the first cycle) and extension at 72°C for 60 seconds and 25 cycles of conventional PCR (done similarly as per touch-down PCR except for the annealing temperature fixed at 57°C).

Amplified mutant Ag 43 signal peptide gene libraries were analysed with gel electrophoresis and GelApp (Sim et al., 2015) and gel extracted. Error rates were estimated by calculating the PCR duplications required to achieve the final gel extracted products yield from 2.5 ng of template, following the GeneMorph II epPCR kit manufacturer’s suggestions. The mutant Ag43 signal peptide libraries were used as megaprimers for the circular PCR amplification using the wildtype pCDF_Ag43sp_2xstrepSfGFPβLac as a template. Briefly, circular PCR reactions were set up with the Platinum SuperFi II Green PCR master mix; the reaction contained 50 ng of Plasmid Template and 250 ng of mutagenized Ag43 signal peptide mega primer libraries. The thermocycling profile is as follows: initial denaturation at 98°C for 120 seconds, followed by 25 cycles of denaturation at 98°C for 30 seconds, annealing at 60°C for 30 seconds and extension at 72°C for 210 seconds. Mutant plasmid libraries were then subjected to a PCR clean-up, eluted with 40 μl nuclease-free water, and treated with 20 U of DpnI (NEB, Singapore; Cat: R0176L) for 30 minutes according to the manufacturer’s instructions.

### Expression of Autotransporter system

To generate mutant clones for expression and selection utilising FACs, ~ 100 ng of Ag43 signal peptide mega primer libraries PCR amplified plasmids were transformed into 100 μl vials of competent BL21 (DE3) cells (Chan et al., 2013).

Transformed cells were then divided and plated onto three plates (Biomedia, Singapore; 100 x 15 mm; Cat: BMH.990000PQ-20) of LB agar containing the appropriate antibiotic and incubated at 37 °C overnight. Colonies were counted with the aid of the APD Colony Counter App (Wong et al., 2019). Transformed cells were collected each round and pooled into LB containing 200μg / ml of spectinomycin. To preserve the diversity of the mutant libraries, induction was performed immediately by diluting the pooled cultures to OD_600_ 0.4 in 5 ml of LB containing 0.5 mM of IPTG and appropriate antibiotics to be incubated at 25°C, 250 rpm (in a Benchtop ES-20, Orbital Shaker-Incubator, Latvia) for 5 to 6 hours. The controls: untransformed (no Plasmid or NoP) and wild-type plasmid containing BL21 (DE3), were induced in the same manner with no antibiotics included for the untransformed cells.

### Cell sorting and Flow Cytometry

For flow cytometric assessments of the heterologous display of sfGFP on mutant clones, 500 μl of cells were first normalized to OD_600_ 0.4. Next, cells were washed twofold with 500μl of a phosphate-buffered solution containing protease inhibitor (PI PBS). Next, staining was performed with 1.5 μg/ml of streptavidin conjugated to Alexa Fluor™ 647 (Invitrogen ™, Singapore) for an hour, followed by two additional washes with PI PBS before acquisition and analysis on the CytoFlex Srt (Beckman Coulter, Singapore; Model: V5-B2-Y5-R3). Stained cells were then stored at 4 °C for up to 24 hours. Cell sorting was performed at an OD_600_ of 0.2.

### Flow cytometric and FACs assay setup

Streptavidin-bound cells were detected in APC (660/20 bandpass filter) herein called 2xstrep-APC, and sfGFP displayed on the cell surface was detected in FITC (525/40 bandpass filter) herein called sfGFP-FITC, while the control, BL21 (DE3) was set for baseline adjustment. After 5 million events were collected, dual colour flow cytometric density plots were generated. Next, 2xstrep-APC+ and sfGFP-FITC+ cells were sorted (Threshold for sort-gate: 2xstrep-APC ~3×10^4^ to 5×10^5^, sfGFP-FITC intensity ~ 4×10^4^ to 3×10^6^) in purity mode into FACs tubes between 3 rounds of mutagenesis and FACs selection of Ag43 signal peptide mutant libraries. In the first two rounds, cells were tube sorted. Next, cells were mixed with 20 ml of media split between two 50 ml Falcon™ tubes. In the last round, cells from round 2 were induced the following day and FACs enriched for 2xstrep-APC+ and sfGFP-FITC+ clones. Clones were then miniprepped, subjected to a round of mutagenesis, allowed to express and index sorted into 96 well microtiter plates at the end of the third round of sorting. Index sorted cells were transferred to deep well plates containing 400μl of media. Cells in either tubes or deep well plates were incubated at 37 °C, 250 rpm overnight in the ES-20, Orbital Shaker-Incubator and MaxQ 8000 incubator (ThermoFisher, U.S.A), respectively.

### Screening of the mutant clones

For all experiments that involved the expression of clones for screening and characterization, the MaxQ 8000 incubator was used. Overnight cultures of 43 mutant clones isolated with the CytoFlex Index sorting function were inoculated as 1% starter culture in 2 ml LB medium (in 14 ml flacon snap cap tubes; Corning, USA) induced at an OD_600_ of 0.6 to 0.8. Cells were expressed with 0.5 mM IPTG at 25°C, 250 rpm for 5 to 6 hours.

To characterize the total sfGFP displayed, 1% starter cultures were inoculated in 10 ml of LB (in 50 ml falcon tubes; Corning, USA) and induced similarly as were performed for initial screens. After 6 and 12 hours of induction, 5 ml of culture were collected at the time points. To determine total fluorescent sfGFP on the surface of cells, the bacteria cells were normalized by OD where they were first spun down and resuspended to the equivalent of OD 4 or OD 8 with 400μl of 2.5g/L of Trypsin solution (Nacali Tesque, Japan; Cat: 32777-44, Lot No. L0F24273), followed by incubation at 37 °C, 250 rpm for 2 hours. The supernatant of trypsinized cells sfGFP was separated by centrifugation while the cell pellet was resuspended with the same volume of fresh trypsin solution.

For cells trypsinized at OD 4, 150 μl of trypsinised cell supernatant or trypsinized cell pellet resuspended in fresh trypsin solution were analysed for sfGFP fluorescence (485/20 nm, emission 528/20 nm; optimized gain manually set at 75) with a TECAN SAFIRE II plate-reader (TECAN, Switzerland). Cleaved sfGFP from the supernatant of cells trypsinized at OD 8 (gave bright bands from initial optimizations on bacteria cells, including the wild-type signal peptide) were used for SDS-PAGE separation of functional sfGFP. The cleaved sfGFP-containing liquid was mixed with 5x native loading dye(Tris-HCl 250 mM, pH 6.8; 30%, glycerol; 0.02% Bromophenol blue) without a subsequent boiling step and loaded (13ul) using commercially available precast 12% SDS-PAGE gels (BioRad, Singapore; Cat: 4561046). Gels were illuminated with a blue light transilluminator. ImageLab™ (BioRad, USA) and GelApp (Sim et al., 2015) were used for densitometric analysis and band analysis, respectively.

To sequence the mutants, Sanger sequencing (1^st^ Base, Singapore) was performed on PCR amplification using the Platinum SuperFi II Green PCR master mix (following the manufacturer’s protocol, using primer set: Ag43SpSeq_F, 5’- ATC GAT GTC TCG ATC ACG TCG CGG GAA TTG TGA GCG GAT AAC AAT −3’; and Ag43SpSeq_R 5’- GCA CCA GAC GGT TGC CAC AGG CAT CTT TGC TCA SGCA CGC TTT GGG −3’) as the quality of BL21 (DE3) isolated plasmids were poor. Sequences were analysed with a locally installed version of YAQAAT (Koh et al., 2021) (using the wildtype signal peptide sequence as a template) and DNAApp (Nguyen et al., 2014) for analysis.

### Data analysis

The plate-reader and densitometric data were analysed and plotted using GraphPad Prism Version 9.0.0; a One-way Analysis of Variance (ANOVA) with Dunnett’s comparison test against the induced wildtype was performed. FACs and flow-cytometric plots were generated with the CytExpert SRT version 1.0.2 and FlowJo™ version 10.8.1 software.

## RESULTS AND DISCUSSION

### FACs selection for single-cell mutants with high levels of display and expression

The native *E. coli* Ag43 autotransporter displays heterologous proteins on the bacteria cell surface and is a good marker for our FACS-sorted high throughput selection of single-cells. To detect extracellular display, fluorescent-tagged streptavidin and 2xstrep-tag fused to a reporter of protein expression – sfGFP - was used. Three rounds of selection (denoted as R1,2, and 3) of reiterative mutagenesis and selection for single-cell mutants with extreme upper right quadrant sfGFP-FITC and 2xstrep-APC signals screening of 2900 – 8700 colonies showed evident enrichment. The average error rates of mutant libraries were estimated to be between medium to high (4.5–16 average errors / 1000 bp) range across all the three rounds of mutagenesis.

To compare the various rounds, selected clones were glycerol stocked, thawed, expressed, and subjected to flow cytometry (Figure 2A). We noticed the 2xstrep-APC+/sfGFP-FITC+ (positive for APC. and FITC) wild-type to exhibit a wave-tide-like topology (Figure 2A, B) around 1×10^4^ AFUs on the flow cytometry plots.

**Figure 2.**
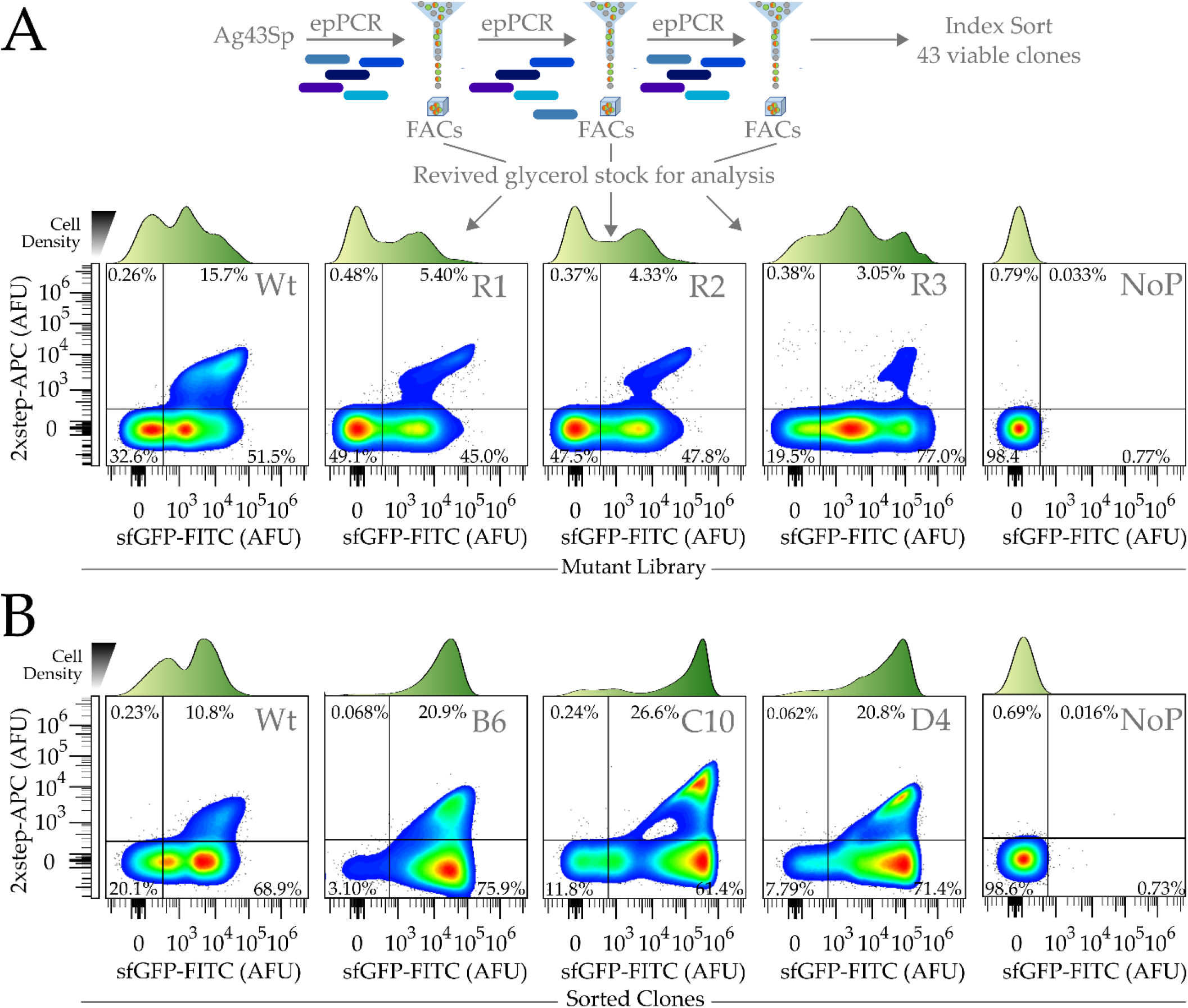
Representative flow cytometric charts of mutant Ag43 signal peptide BL21 libraries. (A) Flow cytometry of sorted mutant Ag43 signal peptide libraries. R1, R2, and R3 corresponded to cell populations with mutant libraries that underwent 1,2 and 3 rounds of epPCR mutagenesis and FACs selection. Wt and NoP corresponded to populations containing Wildtype Ag43 signal peptide and no plasmid, respectively. The population of single-cells 2xstrep-APC+/sfGFP-FITC+ (which corresponded to higher fluorescent streptavidin binding and sfGFP fluorescence) and those that exhibited extreme sfGFP-FITC signals were found reduced and increased, respectively, over the three rounds of selection. To express the displayed proteins, 100 μl of frozen bacteria glycerol stocks were used as a starter culture to inoculate 10 ml cultures in Falcon tubes (B)Flow cytometry of the top 3 mutants (B6, C10 and D4) out of 43 screened clones with the AFU, arbitrary fluorescence units. Histograms of the positive *E. coli* for different levels of sfGFP-FITC fluorescence are shown above the respective plots.

We observed the mutant 2xstrep-APC+/sfGFP-FITC+ and 2xstrep-APC-/sfGFP-FITC+ sub-populations to progressively increase in sfGFP-FITC signals over the three rounds of mutagenesis and selection (Figure 2A). Single-cells within R1 and R2 mutant pools positive for 2xstrep-APC+ formed a sizable population with signals around ~1×10^5^ AFUs compared to ~ 1×10^4^ AFUs in the wildtype, demonstrating the successful selection pressure applied for cells with stronger sfGFP signals (Figure 2A). Between R1 and R2, the increased single-cell sfGFP-FITC fluorescence was accompanied by a corresponding decrease in 2xstrep-APC+/sfGFP-FITC+ single-cell populations. This could be due to the number of cells expressing 2xstrep-APC signals for surface display being decreased (Figure 2A) or fluorescence quenching that could be seen by the boomerang-shaped dot plots of the 2xstrep-APC+/sfGFP-FITC+.

By the third round (R3), there was a significant decrease of 2xstrep-APC+/sfGFP-FITC+ populations that was accompanied by major populations that were either 2xstrep-APC+ or 2xstrep-APC- to express sfGFP-FITC around 1×10^5^ AFUs which was 2-10-fold higher than the majority in the wildtype control. Given the inverse relationship between 2xstrep-APC+ and sfGFP-FITC+ signals, there was an unexplained tradeoff in the selection.

In the final third round of selection, we isolated 43 viable clones sorted into 96 well plates (Figure 2B, Figure S1-3), identifying three representative mutant clones: B6, C10 and D4, which displayed sfGFP-FITC+/2xstrep-APC+ population enrichments of 20.9%, 26.6% and 20.8% the initial screens, respectively (Figure 2B). Within the population of sfGFP-FITC+/2xstrep- APC+ and sfGFP-FITC+/2xstrep-APC-, a sizable population of B6, C10 and D4 cells displayed sfGFP-FITC fluorescence around 3×10^4^, 2×10^5^, and 1×10^5^ AFUs respectively. In contrast, most of the cells in the wild-type population were around 1×10^4^ AFUs for sfGFP-FITC fluorescence.

### Characterization of bulk sfGFP expression in mutants

As single cell population data may not translate to bulk titers of displayed sfGFP and to further confirm that sfGFP were indeed displayed on the surface, we trypsinized untransformed BL21 (DE3), wild-type (Induced and uninduced) and mutant clones (C10, D4, B6) Ag43 leader signal peptide constructs that were induced for 6 and 12 hours respectively. Since sfGFP is resistant to trypsinization(Chiang et al., 2001), it would remain intact and be released while the bacterial cell walls remain intact. By analyzing the supernatant with a plate-reader and native SDS-PAGE (boiling omitted to preserve sfGFP functionality), we would be able to verify that the protein was released from the bacterial surface. To quantify the density of bands in native SDS-PAGE across different gels, the first induced wildtype replicate (Wt I-R1) was used as an internal control for densitometric analysis (Figure 3B).

**Figure 3.**
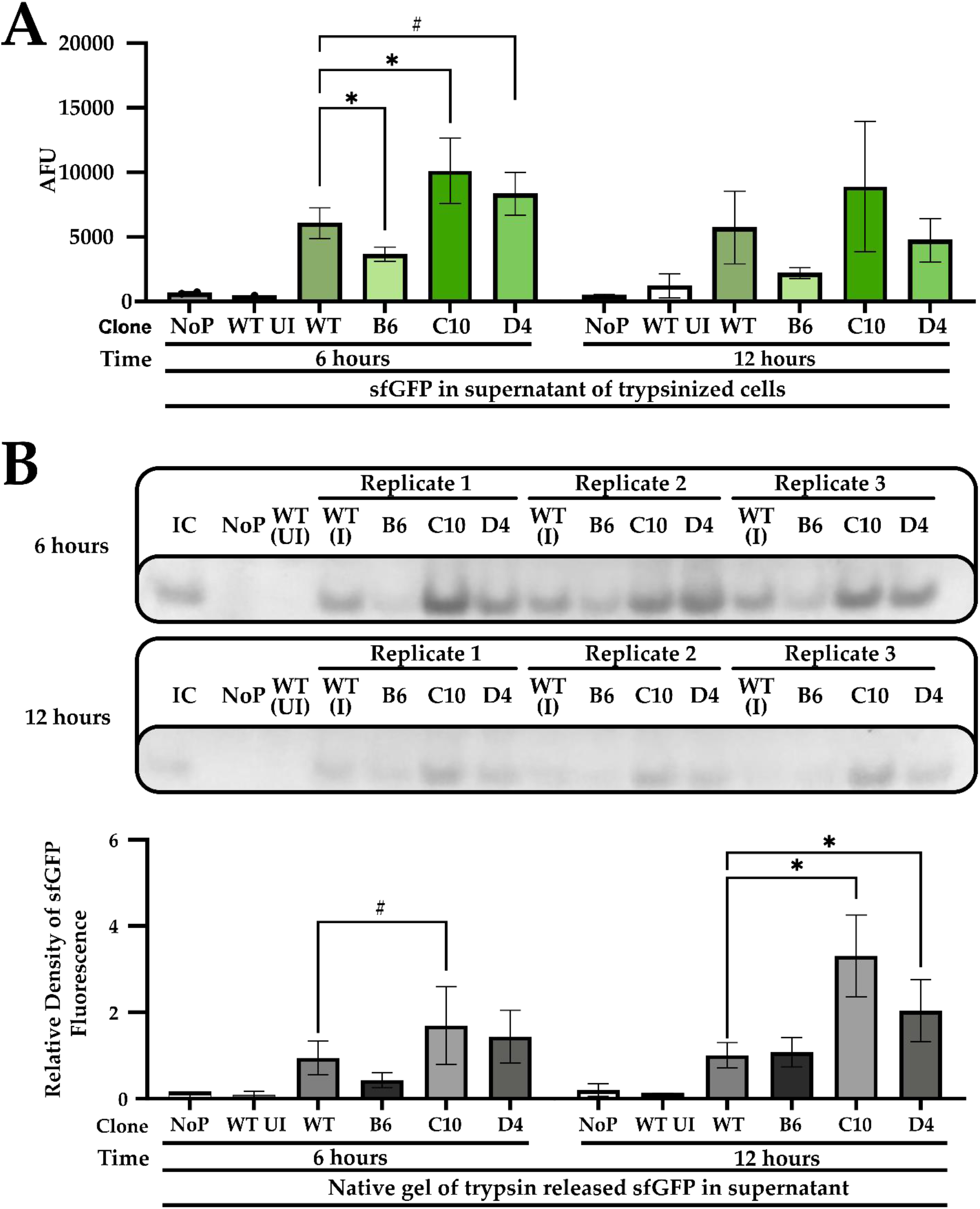
Quantification and SDS PAGE of trypsinized bacterial cells to trypsinized sfGFP in the supernatant. (A) Mean total sfGFP fluorescence of cell supernatants after trypsinization. Bacterial cells of the wildtype (Wt, induced) Ag43 signal peptide was compared with mutant (B6, C10 and D4) Ag43 variants. Uninduced wildtype Ag43 signal peptide (Wt UI) and BL21 (DE3) untransformed or with no plasmid (NoP) served as negative controls. (B) Native SDS-PAGE gel of trypsinized bacterial cell supernatants. sfGFP were excited by the blue light. A common internal control(IC) corresponding to Replicate 1 of WT (I) was included to facilitate comparison between gels. (C) Densitometric analysis of sfGFP signals is shown in (B). Except for Wt UI and NoP, WT and mutant variants were performed in triplicates and sextuplicate, respectively. One-way ANOVA and Dunnett posthoc test were used for comparing the wildtype and mutant variants (#<0.1,*,p<0.05).

Examining trypsinzed bacteria cell supernatants at the 6 and 12-hour timepoints by plate-reader microtiter assays showed that at the 6-hour time point, sfGFP fluorescence of C10 and D4 were significantly higher by 1.7 (p<0.05), and 1.4 (p<0.1) folds than the induced wildtype controls (Wt). On the contrary, B6 had lower fluorescence than the wildtype with similar trends observed for the 12-hour timepoint experiments for all the three mutants with B6, C10 and D4 but with readings of 0.4 and 1.6 and 0.8 folds to the wildtype, and without statistical significance. Considering that the readings from the supernatant also showed higher cleaved sfGFP levels, the proteins were displayed on the cell surface.

The results from the Native SDS-PAGE were congruent with the trends found in the plate-reader-based microtiter assay. At the 6-hour time point, densities of the fluorescent signal generated by sfGFP separated by native SDS-PAGE showed B6, D4 and C10 to be 0.5, 1.8 and 1.5 folds higher than the wild-type (Wt in Figure 3B) with only the change in C10 to be significant(p<0.1). At the 12-hour time point, sfGFP fluorescence densities of C10 and D4 were found to be significantly increased by 3.2 and 2-fold. B6 was, however, unchanged compared to Wt (Figure 3B, p<0.05).

Observing trypsinized cells to glow strongly under blue light and populations of 2xstrep-APC-cells to be sfGFP-FITC^(+)^, we also sought to quantify the intracellular fluorescence of post-trypsinized whole cells (with a plate reader, Figure 4A). Intracellular fluorescence of all mutants were significantly higher than the wild-type (p<0.05) (Figure 4B), with B6, C10 and D4 clones exhibiting 4.2, 26.6 and 8.9 folds than the wildtype at the 6 hours timepoint. At the 12 hours, point B6, C10, and D4 clones showed 7.7, 25.8 and 13-folds more fluorescence than the wild-type. The data suggest that the mutant Ag43 signal peptides were more efficient than the wild-type after the three rounds of selections. Intriguingly the observed boosts in intracellular sfGFP accumulation were asymmetric relative to the modest improvements in the display of the sfGFP-beta-lactamase fusion. This hinted at the greater complexity of signal peptide-directed protein export pathways, particularly for the fast-folding sfGFP in *E. coli* (Dinh and Bernhardt, 2011; Fisher and DeLisa, 2008).

**Figure 4.**
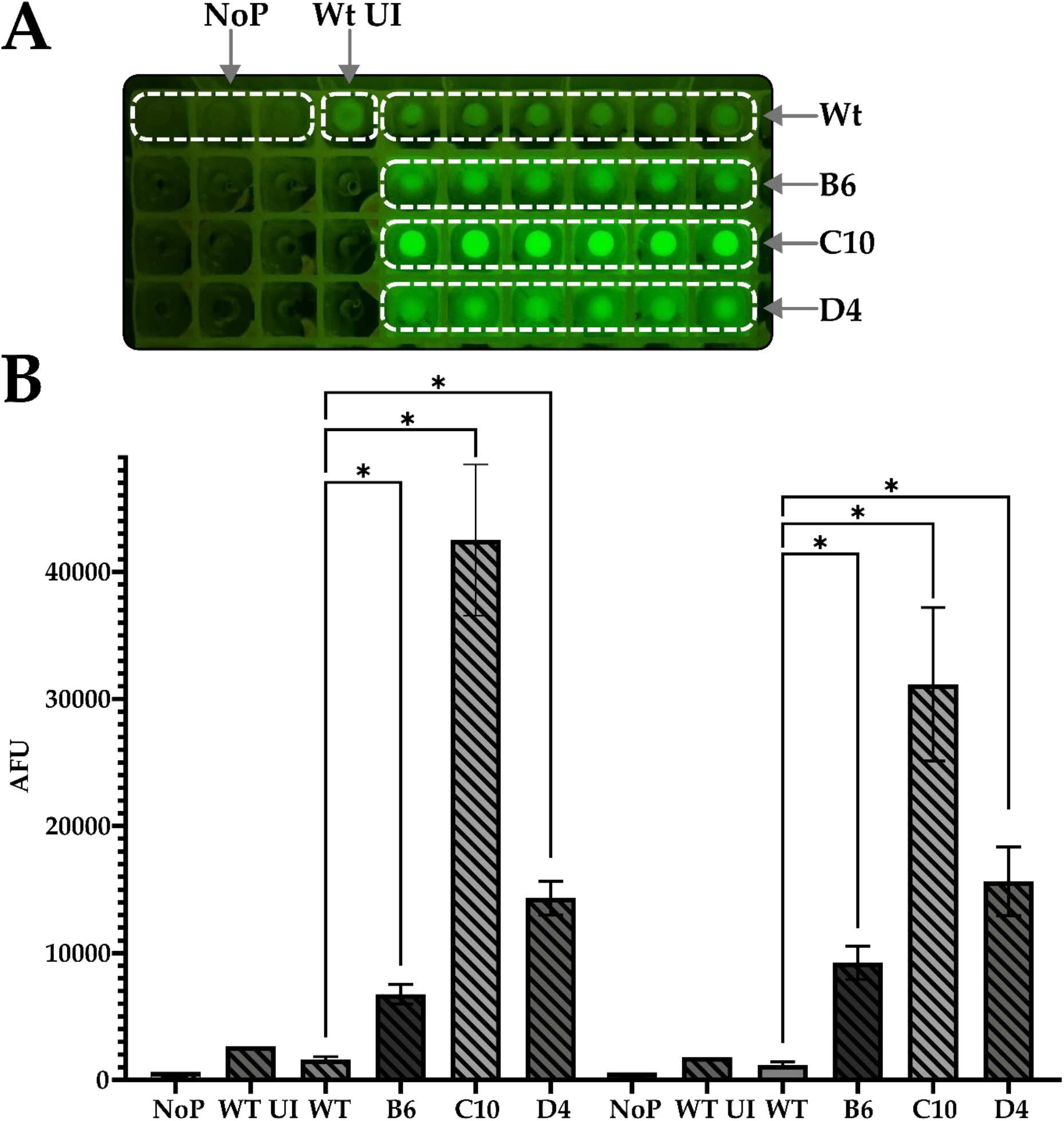
Quantification of whole bacterial cell sfGFP. (A) A photograph of pelleted trypsinized whole bacterial cells (from figure 3) expressing sfGFP fused to either Wild-type (Wt, induced) or mutant (B6, C10 and D4) Ag43 sp, Pelleted cells with uninduced SfGFP fused to wild-type Ag43 Sp (Wt UI) or BL21 (De3) without plasmid (NoP) served as negative controls. (B) Intracellular sfGFP signals from pelleted cells (from A, resuspended in trypsin solution) were quantified with a microplate reader. Except for Wt UI and NoP, WT and mutant variants were performed in triplicates and sextuplicates, respectively. One-way ANOVA and Dunnett posthoc test were used to compare the wild-type and selected mutant variants (*,p<0.05).

Selected mutants from seven other picked clones were sequenced and found to converge on the sequences of both C10 and D10 (Figure 3) which consistently gave higher readings in both whole cells and supernatants. B6 was omitted as its FACS readings as 2xstrep-APC+ populations were around 3×10^4^, AFUs and further analysis of the cleaved sfGFP supernatant readings (fluorescence readings of B6 were 0.6 and 0.4 folds compared to wild-type) were not significantly higher than the wild-type, suggesting no improved surface transport.

Analysis of the sequences showed Clone C10 to have five missense nucleotide mutations 70G>A, 74T>G, 86G>C, 104G>A, 116T>A, and 121C>G leading to amino acid substitutions of A24T, S25A, R29P, G35D, V39D and L41V (Figure 5). Three other clones were also sequenced to harbour the same mutations as C10: B2, C2 and B11 (Table 1, Figure S1-2). On the other hand, Clone D4 had just one silent mutation, G156C (Figure 5), and carried the same mutation as Clone B9 (Table 1, Figure S2). The 156G>C mutation led to a change from a high (CTG) to low usage codon (CTC) for *E. coli* (using strain *E. coli O157:H7 str. EDL933* as a reference) with frequencies of 51.1% and 10.5% respectively (Nakamura, 2022; Nakamura et al., 2000). The presence of silent mutations leading to codons with lower usages was previously reported to increase the ability of protein secretion tags to export proteins extracellularly by 1.6 folds (Gonzalez-Perez et al., 2021) and was recently reported to affect mutational fitness in eukaryotes (Shen et al., 2022).

**Figure 5.**
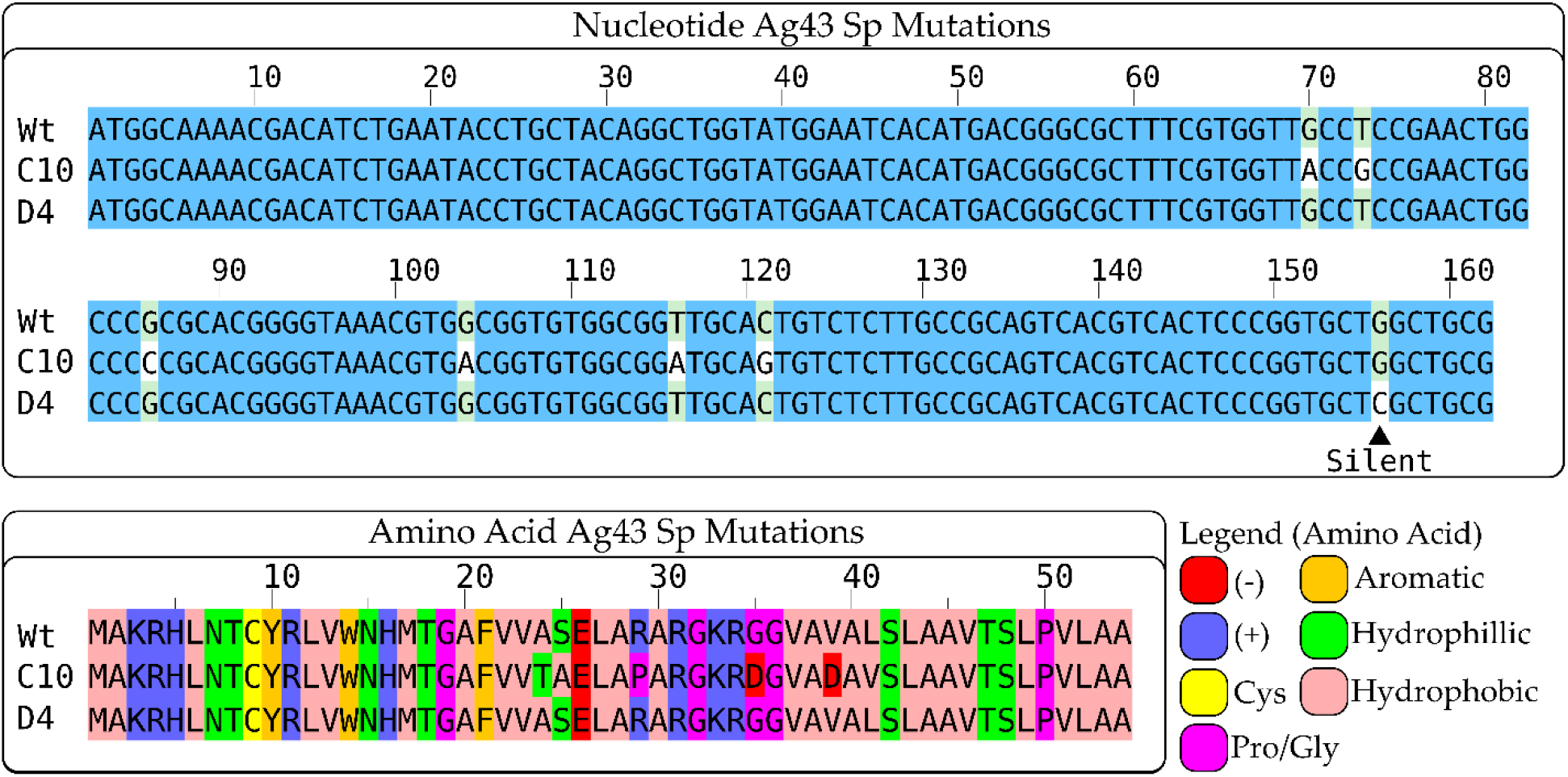
Multiple sequence alignment of the nucleotide and amino acid sequence of the Ag43 signal peptides of Clone C10 and D4; ▲ indicates the position of a silent mutation(Clone D4, G156C). Aligned amino acid sequences are coloured according to the Zappo scheme. The figure was prepared with Jalview(Waterhouse et al., 2009).

**Table 1.**
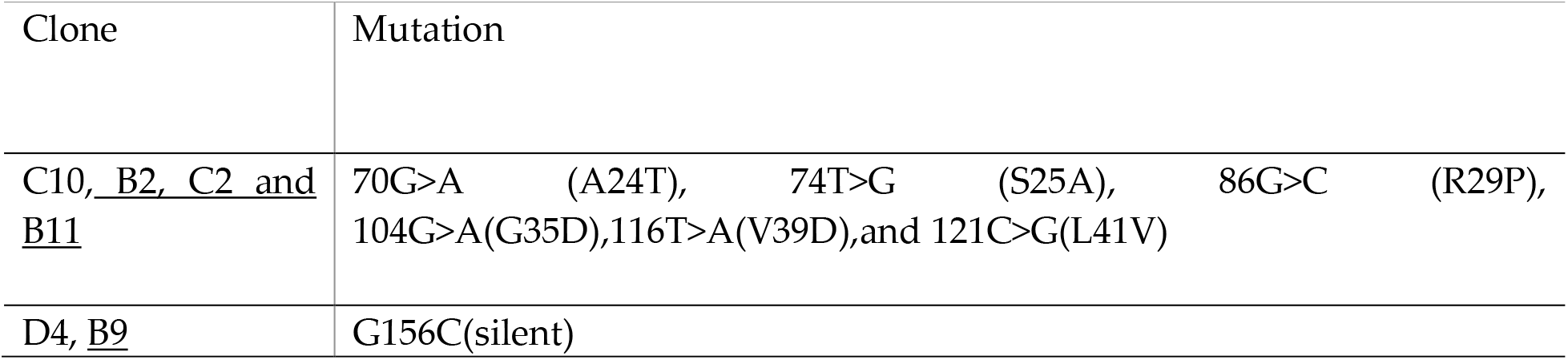
Mutations in sequenced mutant clones, underlined clone indicates mutant clones that shared the same signal peptide mutations as clones C10 and D4.

The use of FACs and flow cytometric screening has been widely cited as a foundational tool for cell selection and is multiparametric and high through-put to perform the selection of biological molecules with novel properties and functions. In this study, we were able to mutate and select signal peptides of autotransporters by relying on two parameters, the expression of sfGFP measured by 2xStrep-APC and total sfGFP (detected as sfGFP-FITC) to generate mutations that facilitated the ability of 2xStrep-APC+ single-cells to exhibit progressively greater sfGFP-FITC fluorescence over three rounds of epPCR mutagenesis and FACs screenings. In this proof-of-concept methodology, we demonstrated our workflow to generate Clones C10 and D4, which harboured five missense and one silent mutation, respectively, for the augmented display of the fast-folding sfGFP-beta-lactamase fusion as measured by plate-reader based analysis and native SDS-PAGE of trypsinized surface proteins.

Employing FACs to screen for mutant epPCR generated mutant Ag43 signal peptide libraries, we were able to screen upwards of 10^5^ bacteria over three rounds of selection. However, this could be scaled up easily as it is only limited by the plating and transformation steps and the throughput of the FACs machine, - which can record a million events in less than two minutes. We anticipate that further development and deployment of this methodology to bio foundries in a highly automated(Chory et al., 2021; DeBenedictis et al., 2022; Pham et al., 2017) and multiplexed fashion could lead to scalable and simultaneous optimization signal peptides, chaperones and autotransporters that direct protein export for different proteins simultaneously. Further development of such methods for optimizing autotransporter-based display systems would contribute to the facile engineering of ELMs, whole cell catalysts, and living medicines (by optimizing the display of enzymes, binders, and functional biologics).

In conclusion, we demonstrated that single-cell bacterial cell sorting coupled with epPCR and the analysis of trypsinized supernatants is a suitable workflow to select for Ag43 signal peptides that could augment bacterial display. The methodology could be adapted for other proteins that could be useful for improving the efficiency of bacterial autotransporter systems for display and protein export.

## Author Contributions

Conceptualization, DWS. Koh, J.H.T and S.K-E.G. Methodology, D.W.-S.K and S.K-E.G. Formal analysis, DWS. Koh, J.H.T and S.K-E.G. Investigation, DWS. Koh and JHT Validation, DWS. Koh and JHT Writing—Original Draft, DWS. Koh, J.H.T and S.K-E.G. Writing—Review & Editing, DWS. Koh, J.H.T and S.K-E.G. Funding Acquisition, S.K-E.G. Supervision, S.K-E.G.

## Acknowledgement

We thank JYY for valuable suggestions and the performing of replicate experiments, AMC for assistance on the FACS and molecular biology, CHKN for programming the OT2 and assistance in some of the experiments, and ZSLH for valuable discussions.

---

**Figure S1.**
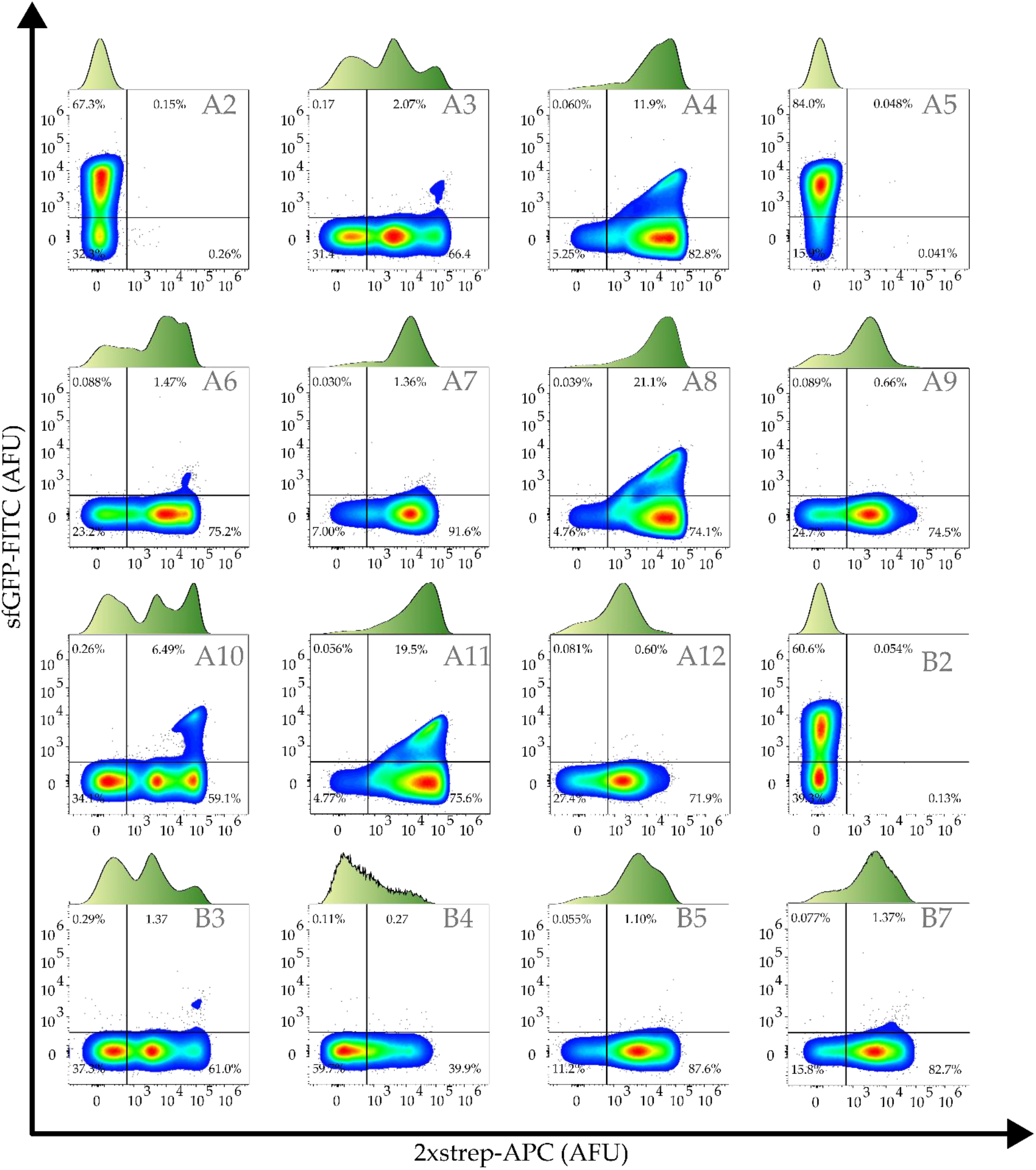
Representative flow cytometric analysis of screened clones (Labelled as A2-D12)

**Figure S2.**
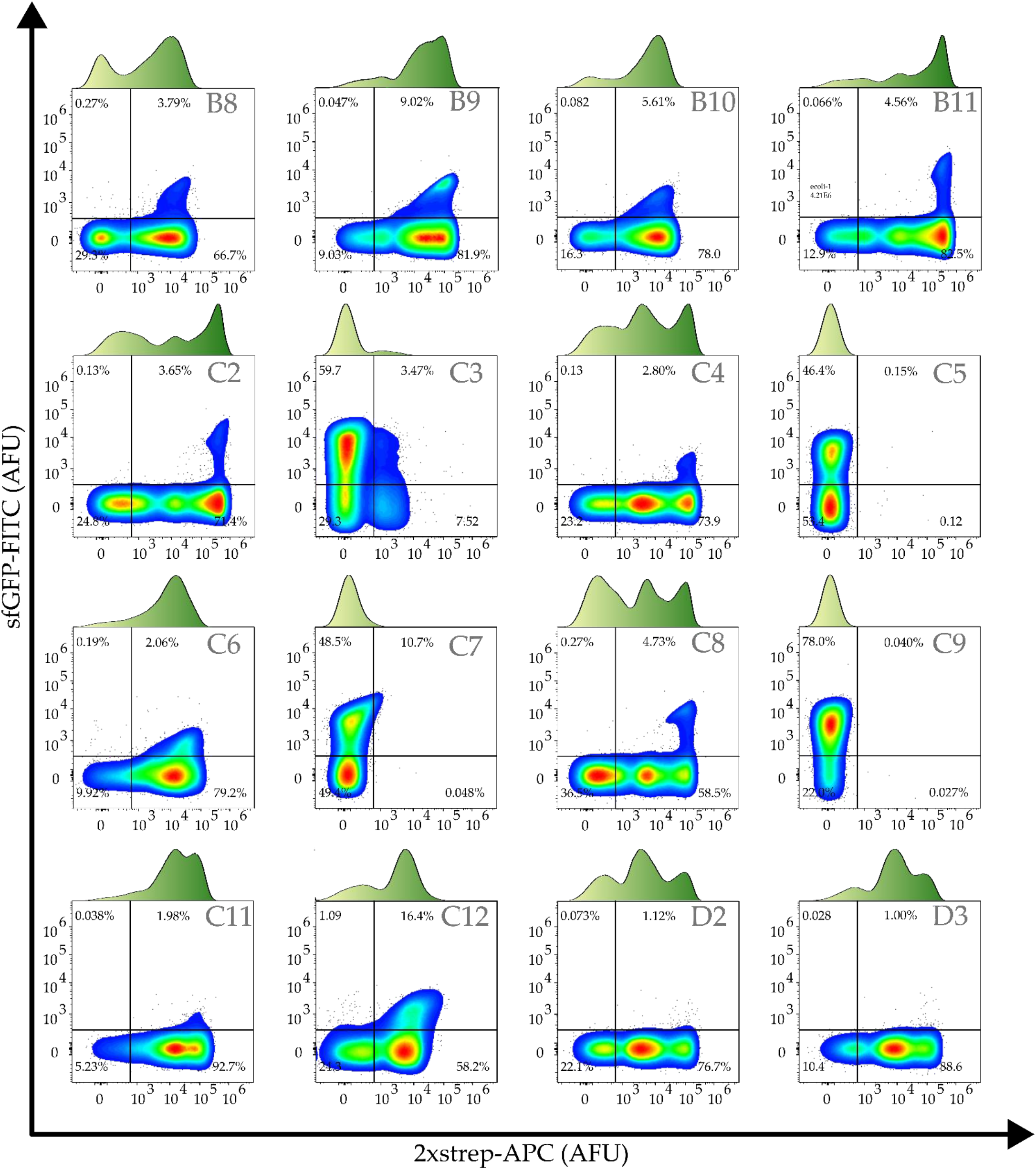
Representative flow cytometric analysis of screened clones (Labelled as A2-D12)

**Figure S3.**
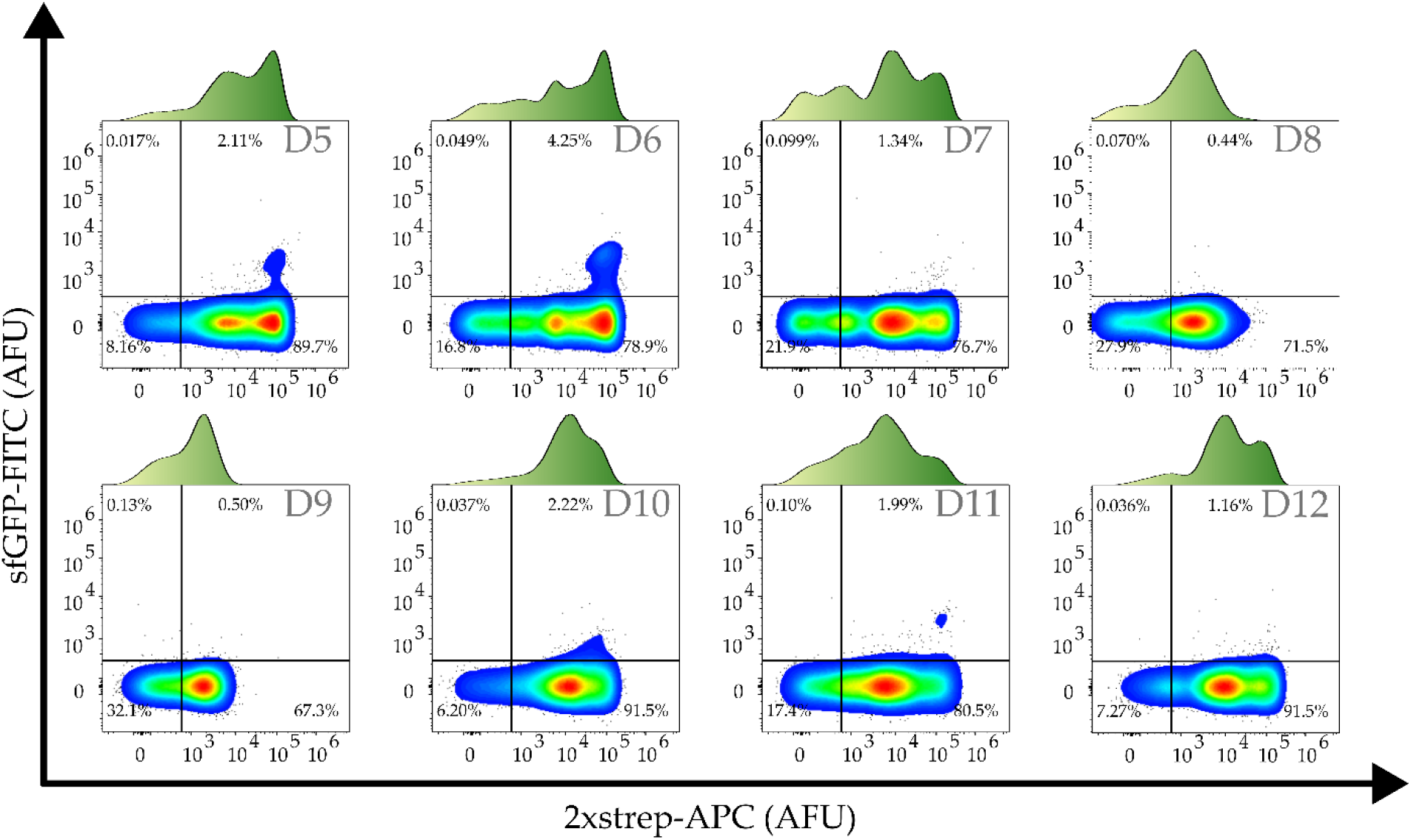
Representative flow cytometric analysis of screened clones (Labelled as A2-D12)

## References

Ahan, R.E., Kirpat, B.M., Saltepe, B., Şeker, U.Ö.Ş., 2019. A Self-Actuated Cellular Protein Delivery Machine. ACS Synth. Biol. 8, 686–696. https://doi.org/10.1021/acssynbio.9b00062

Alessa, A.H.A., Tee, K.L., Gonzalez-Perez, D., Omar Ali, H.E.M., Evans, C.A., Trevaskis, A., Xu, J.-H., Wong, T.S., 2019. Accelerated directed evolution of dye-decolorizing peroxidase using a bacterial extracellular protein secretion system (BENNY). Bioresources and Bioprocessing 6, 20. https://doi.org/10.1186/s40643-019-0255-7

Azam, A., Li, C., Metcalf, K.J., Tullman-Ercek, D., 2016. Type III secretion as a generalizable strategy for the production of full-length biopolymer-forming proteins. Biotechnology and Bioengineering 113, 2313–2320. https://doi.org/10.1002/bit.25656

Beal, J., 2015. Bridging the Gap: A Roadmap to Breaking the Biological Design Barrier. Frontiers in Bioengineering and Biotechnology 2.

Burdette, L.A., Wong, H.T., Tullman-Ercek, D., 2021. An optimized growth medium for increased recombinant protein secretion titer via the type III secretion system. Microbial Cell Factories 20, 44. https://doi.org/10.1186/s12934-021-01536-z

Chan, W.-T., Verma, C.S., Lane, D.P., Gan, S.K.-E., 2013. A comparison and optimization of methods and factors affecting the transformation of Escherichia coli. Bioscience Reports 33, e00086. https://doi.org/10.1042/BSR20130098

Chiang, C.F., Okou, D.T., Griffin, T.B., Verret, C.R., Williams, M.N., 2001. Green fluorescent protein rendered susceptible to proteolysis: positions for protease-sensitive insertions. Arch Biochem Biophys 394, 229–235. https://doi.org/10.1006/abbi.2001.2537

Chory, E.J., Gretton, D.W., DeBenedictis, E.A., Esvelt, K.M., 2021. Enabling high-throughput biology with flexible open-source automation. Mol Syst Biol 17, e9942. https://doi.org/10.15252/msb.20209942

Cui, M., Sun, T., Li, Shubin, Pan, H., Liu, J., Zhang, X., Li, L., Li, Shanshan, Wei, C., Yu, C., Yang, C., Ma, N., Ma, B., Lu, S., Chang, J., Zhang, W., Wang, H., 2021. NIR light-responsive bacteria with live bio-glue coatings for precise colonization in the gut. Cell Reports 36, 109690. https://doi.org/10.1016/j.celrep.2021.109690

Daugherty, P.S., Chen, G., Iverson, B.L., Georgiou, G., 2000. Quantitative analysis of the effect of the mutation frequency on the affinity maturation of single chain Fv antibodies. Proceedings of the National Academy of Sciences 97, 2029–2034. https://doi.org/10.1073/pnas.030527597

DeBenedictis, E.A., Chory, E.J., Gretton, D.W., Wang, B., Golas, S., Esvelt, K.M., 2022. Systematic molecular evolution enables robust biomolecule discovery. Nat Methods 19, 55–64. https://doi.org/10.1038/s41592-021-01348-4

DeLisa, M.P., Samuelson, P., Palmer, T., Georgiou, G., 2002. Genetic Analysis of the Twin Arginine Translocator Secretion Pathway in Bacteria *. Journal of Biological Chemistry 277, 29825–29831. https://doi.org/10.1074/jbc.M201956200

Dinh, T., Bernhardt, T.G., 2011. Using Superfolder Green Fluorescent Protein for Periplasmic Protein Localization Studies ▿. J Bacteriol 193, 4984–4987. https://doi.org/10.1128/JB.00315-11

Duraj-Thatte, A.M., Manjula-Basavanna, A., Rutledge, J., Xia, J., Hassan, S., Sourlis, A., Rubio, A.G., Lesha, A., Zenkl, M., Kan, A., Weitz, D.A., Zhang, Y.S., Joshi, N.S., 2021. Programmable microbial ink for 3D printing of living materials produced from genetically engineered protein nanofibers. Nat Commun 12, 6600. https://doi.org/10.1038/s41467-021-26791-x

Fisher, A.C., DeLisa, M.P., 2008. Laboratory Evolution of Fast-Folding Green Fluorescent Protein Using Secretory Pathway Quality Control. PLoS One 3, e2351. https://doi.org/10.1371/journal.pone.0002351

Georgiou, G., 2001. Analysis of large libraries of protein mutants using flow cytometry, in Advances in Protein Chemistry, Evolutionary Protein Design. Academic Press, pp. 293–315. https://doi.org/10.1016/S0065-3233(01)55007-X

Glass, D.S., Riedel-Kruse, I.H., 2018. A Synthetic Bacterial Cell-Cell Adhesion Toolbox for Programming Multicellular Morphologies and Patterns. Cell 174, 649–658.e16. https://doi.org/10.1016/j.cell.2018.06.041

Gonzalez-Perez, D., Ratcliffe, J., Tan, S.K., Wong, M.C.M., Yee, Y.P., Nyabadza, N., Xu, J.-H., Wong, T.S., Tee, K.L., 2021. Random and combinatorial mutagenesis for improved total production of secretory target protein in Escherichia coli. Sci Rep 11, 5290. https://doi.org/10.1038/s41598-021-84859-6

Griffiths, A.D., Tawfik, D.S., 2003. Directed evolution of an extremely fast phosphotriesterase by in vitro compartmentalization. The EMBO Journal 22, 24–35. https://doi.org/10.1093/emboj/cdg014

Gustavsson, M., Bäcklund, E., Larsson, G., 2011a. Optimisation of surface expression using the AIDA autotransporter. Microbial Cell Factories 10, 72. https://doi.org/10.1186/1475-2859-10-72

Gustavsson, M., Bäcklund, E., Larsson, G., 2011b. Optimisation of surface expression using the AIDA autotransporter. Microbial Cell Factories 10, 72. https://doi.org/10.1186/1475-2859-10-72

Haitjema, C.H., Boock, J.T., Natarajan, A., Dominguez, M.A., Gardner, J.G., Keating, D.H., Withers, S.T., DeLisa, M.P., 2014. Universal Genetic Assay for Engineering Extracellular Protein Expression. ACS Synth. Biol. 3, 74–82. https://doi.org/10.1021/sb400142b

Heng, Z.S.-L., Yeo, J.Y., Koh, D.W.-S., Gan, S.K.-E., Ling, W.-L., 2022. Augmenting recombinant antibody production in HEK293E cells: optimizing transfection and culture parameters. Antibody Therapeutics 5, 30–41. https://doi.org/10.1093/abt/tbac003

Ho, C.L., Tan, H.Q., Chua, K.J., Kang, A., Lim, K.H., Ling, K.L., Yew, W.S., Lee, Y.S., Thiery, J.P., Chang, M.W., 2018. Engineered commensal microbes for diet-mediated colorectal-cancer chemoprevention. Nat Biomed Eng 2, 27–37. https://doi.org/10.1038/s41551-017-0181-y

Holowko, M.B., Frow, E.K., Reid, J.C., Rourke, M., Vickers, C.E., 2020. Building a biofoundry. Synth Biol (Oxf) 6, ysaa026. https://doi.org/10.1093/synbio/ysaa026

Hörnström, D., Larsson, G., van Maris, A.J.A., Gustavsson, M., 2019. Molecular optimization of autotransporter-based tyrosinase surface display. Biochimica et Biophysica Acta (BBA) - Biomembranes 1861, 486–494. https://doi.org/10.1016/j.bbamem.2018.11.012

Jin, X., Riedel-Kruse, I.H., 2018. Biofilm Lithography enables high-resolution cell patterning via optogenetic adhesin expression. Proc Natl Acad Sci U S A 115, 3698–3703. https://doi.org/10.1073/pnas.1720676115

Kjærgaard, K., Hasman, H., Schembri, M.A., Klemm, P., 2002. Antigen 43-Mediated Autotransporter Display, a Versatile Bacterial Cell Surface Presentation System. J Bacteriol 184, 4197–4204. https://doi.org/10.1128/JB.184.15.4197-4204.2002

Kleiner-Grote, G.R.M., Risse, J.M., Friehs, K., 2018. Secretion of recombinant proteins from E. coli. Eng Life Sci 18, 532–550. https://doi.org/10.1002/elsc.201700200

Koh, D.W.-S., Chan, K.-F., Wu, W., Gan, S.K.-E., 2021. Yet Another Quick Assembly, Analysis and Trimming Tool (YAQAAT): A Server for the Automated Assembly and Analysis of Sanger Sequencing Data. Journal of Biomolecular Techniques: JBT. https://doi.org/10.7171/jbt.2021-3202-003

Kylilis, N., Riangrungroj, P., Lai, H.-E., Salema, V., Fernández, L.Á., Stan, G.-B.V., Freemont, P.S., Polizzi, K.M., 2019. Whole-Cell Biosensor with Tunable Limit of Detection Enables Low-Cost Agglutination Assays for Medical Diagnostic Applications. ACS Sens. 4, 370–378. https://doi.org/10.1021/acssensors.8b01163

Lindroos, M., Hörnström, D., Larsson, G., Gustavsson, M., van Maris, A.J.A., 2019. Continuous removal of the model pharmaceutical chloroquine from water using melanin-covered Escherichia coli in a membrane bioreactor. Journal of Hazardous Materials 365, 74–80. https://doi.org/10.1016/j.jhazmat.2018.10.081

Ling, W.-L., Lua, W.-H., Poh, J.-J., Yeo, J.Y., Lane, D.P., Gan, S.K.-E., 2018. Effect of VH–VL Families in Pertuzumab and Trastuzumab Recombinant Production, Her2 and FcγIIA Binding. Frontiers in Immunology 9, 469. https://doi.org/10.3389/fimmu.2018.00469

Ling, W.-L., Su, C.T.-T., Lua, W.-H., Poh, J.-J., Ng, Y.-L., Wipat, A., Gan, S.K.-E., 2020. Essentially Leading Antibody Production: An Investigation of Amino Acids, Myeloma, and Natural V-Region Signal Peptides in Producing Pertuzumab and Trastuzumab Variants. Frontiers in Immunology 11.

Ling, W.-L., Yeo, J.Y., Ng, Y.-L., Wipat, A., Gan, S.K.-E., 2022. More Than Meets the Kappa for Antibody Superantigen Protein L (PpL). Antibodies 11, 14. https://doi.org/10.3390/antib11010014

Meuskens, I., Saragliadis, A., Leo, J.C., Linke, D., 2019. Type V Secretion Systems: An Overview of Passenger Domain Functions. Frontiers in Microbiology 10.

Muñoz-Gutiérrez, I., Moss-Acosta, C., Trujillo-Martinez, B., Gosset, G., Martinez, A., 2014. Ag43-mediated display of a thermostable β-glucosidase in Escherichia coli and its use for simultaneous saccharification and fermentation at high temperatures. Microbial Cell Factories 13, 106. https://doi.org/10.1186/s12934-014-0106-3

Nakamura, Y., 2022. Codon usage table [WWW Document]. URL http://www.kazusa.or.jp/codon/cgi-bin/showcodon.cgi?species=155864 (accessed 7.4.22).

Nakamura, Y., Gojobori, T., Ikemura, T., 2000. Codon usage tabulated from international DNA sequence databases: status for the year 2000. Nucleic Acids Research 28, 292. https://doi.org/10.1093/nar/28.1.292

Natarajan, A., Haitjema, C.H., Lee, R., Boock, J.T., DeLisa, M.P., 2017. An Engineered Survival-Selection Assay for Extracellular Protein Expression Uncovers Hypersecretory Phenotypes in Escherichia coli. ACS Synth Biol 6, 875–883. https://doi.org/10.1021/acssynbio.6b00366

Nguyen, P.-V., Verma, C.S., Gan, S.K.-E., 2014. DNAApp: a mobile application for sequencing data analysis. Bioinformatics 30, 3270–3271. https://doi.org/10.1093/bioinformatics/btu525

Pham, H.L., Wong, A., Chua, N., Teo, W.S., Yew, W.S., Chang, M.W., 2017. Engineering a riboswitch-based genetic platform for the self-directed evolution of acid-tolerant phenotypes. Nat Commun 8, 411. https://doi.org/10.1038/s41467-017-00511-w

Poh, J.J., Gan, S.K.-E., 2014. The Determination of Factors involved in Column-Based Nucleic Acid Extraction and Purification. J Bioprocess Biotech 04. https://doi.org/10.4172/2155-9821.1000157

Ramesh, B., Sendra, V.G., Cirino, P.C., Varadarajan, N., 2012. Single-cell Characterization of Autotransporter-mediated Escherichia coli Surface Display of Disulfide Bond-containing Proteins *. Journal of Biological Chemistry 287, 38580–38589. https://doi.org/10.1074/jbc.M112.388199

Schroeder, J., Brown, N., Kaye, P., Aebischer, T., 2011. Single Dose Novel Salmonella Vaccine Enhances Resistance against Visceralizing L. major and L. donovani Infection in Susceptible BALB/c Mice. PLOS Neglected Tropical Diseases 5, e1406. https://doi.org/10.1371/journal.pntd.0001406

Selas Castiñeiras, T., Williams, S.G., Hitchcock, A., Cole, J.A., Smith, D.C., Overton, T.W., 2018. Development of a generic β-lactamase screening system for improved signal peptides for periplasmic targeting of recombinant proteins in Escherichia coli. Sci Rep 8, 6986. https://doi.org/10.1038/s41598-018-25192-3

Shen, X., Song, S., Li, C., Zhang, J., 2022. Synonymous mutations in representative yeast genes are mostly strongly non-neutral. Nature 606, 725–731. https://doi.org/10.1038/s41586-022-04823-w

Sim, J.-Z., Nguyen, P.-V., Hwee-Kuan, L., Gan, S.K.-E., 2015. GelApp: Mobile gel electrophoresis analyser. Nature Methods. https://doi.org/10.1038/an9643

Starr, T.N., Greaney, A.J., Hilton, S.K., Ellis, D., Crawford, K.H.D., Dingens, A.S., Navarro, M.J., Bowen, J.E., Tortorici, M.A., Walls, A.C., King, N.P., Veesler, D., Bloom, J.D., 2020. Deep Mutational Scanning of SARS-CoV-2 Receptor Binding Domain Reveals Constraints on Folding and ACE2 Binding. Cell 182, 1295–1310.e20. https://doi.org/10.1016/j.cell.2020.08.012

Su, C.T.-T., Lua, W.-H., Ling, W.-L., Gan, S.K.-E., 2018. Allosteric Effects between the Antibody Constant and Variable Regions: A Study of IgA Fc Mutations on Antigen Binding. Antibodies (Basel) 7. https://doi.org/10.3390/antib7020020

Su, C.T.-T., Lua, W.-H., Poh, J.-J., Ling, W.-L., Yeo, J.Y., Gan, S.K.-E., 2021. Molecular Insights of Nickel Binding to Therapeutic Antibodies as a Possible New Antibody Superantigen. Front Immunol 12, 676048. https://doi.org/10.3389/fimmu.2021.676048

Taw, M.N., Li, M., Kim, D., Rocco, M.A., Waraho-Zhmayev, D., DeLisa, M.P., 2021. Engineering a Supersecreting Strain of Escherichia coli by Directed Coevolution of the Multiprotein Tat Translocation Machinery. ACS Synth. Biol. 10, 2947–2958. https://doi.org/10.1021/acssynbio.1c00183

Taylor, N.D., Garruss, A.S., Moretti, R., Chan, S., Arbing, M.A., Cascio, D., Rogers, J.K., Isaacs, F.J., Kosuri, S., Baker, D., Fields, S., Church, G.M., Raman, S., 2016. Engineering an allosteric transcription factor to respond to new ligands. Nat Methods 13, 177–183. https://doi.org/10.1038/nmeth.3696

Townshend, B., Kennedy, A.B., Xiang, J.S., Smolke, C.D., 2015. High-throughput cellular RNA device engineering. Nat Methods 12, 989–994. https://doi.org/10.1038/nmeth.3486

van der Woude, M.W., Henderson, I.R., 2008. Regulation and function of Ag43 (flu). Annu Rev Microbiol 62, 153–169. https://doi.org/10.1146/annurev.micro.62.081307.162938

VanArsdale, E., Hörnström, D., Sjöberg, G., Järbur, I., Pitzer, J., Payne, G.F., van Maris, A.J.A., Bentley, W.E., 2020. A Coculture Based Tyrosine-Tyrosinase Electrochemical Gene Circuit for Connecting Cellular Communication with Electronic Networks. ACS Synth. Biol. 9, 1117–1128. https://doi.org/10.1021/acssynbio.9b00469

Veiga, E., de Lorenzo, V., Fernández, L.A., 2003. Neutralizationof Enteric Coronaviruses with Escherichia coli CellsExpressing Single-Chain Fv-AutotransporterFusions. Journal of Virology 77, 13396–13398. https://doi.org/10.1128/JVI.77.24.13396-13398.2003

Warszawski, S., Katz, A.B., Lipsh, R., Khmelnitsky, L., Nissan, G.B., Javitt, G., Dym, O., Unger, T., Knop, O., Albeck, S., Diskin, R., Fass, D., Sharon, M., Fleishman, S.J., 2019. Optimizing antibody affinity and stability by the automated design of the variable light-heavy chain interfaces. PLOS Computational Biology 15, e1007207. https://doi.org/10.1371/journal.pcbi.1007207

Waterhouse, A.M., Procter, J.B., Martin, D.M.A., Clamp, M., Barton, G.J., 2009. Jalview Version 2—a multiple sequence alignment editor and analysis workbench. Bioinformatics 25, 1189–1191. https://doi.org/10.1093/bioinformatics/btp033

Widmaier, D.M., Tullman-Ercek, D., Mirsky, E.A., Hill, R., Govindarajan, S., Minshull, J., Voigt, C.A., 2009. Engineering the Salmonella type III secretion system to export spider silk monomers. Mol Syst Biol 5, 309. https://doi.org/10.1038/msb.2009.62

Wong, C.-F., Yeo, J.Y., Gan, S.K.-E., 2019. Republication – APD Colony Counter App: Using Watershed Algorithm for improved colony counting. Scientific Phone Apps and Mobile Devices. https://doi.org/10.30943/2019/c23122019

Wu, H.-C., Tsao, C.-Y., Quan, D.N., Cheng, Y., Servinsky, M.D., Carter, K.K., Jee, K.J., Terrell, J.L., Zargar, A., Rubloff, G.W., Payne, G.F., Valdes, J.J., Bentley, W.E., 2013. Autonomous bacterial localization and gene expression based on nearby cell receptor density. Mol Syst Biol 9, 636. https://doi.org/10.1038/msb.2012.71

